# TRPV4 Channels Mediate Bladder Cancer Cell Proliferation, Migration and Chemoresistance

**DOI:** 10.1101/2025.06.05.657842

**Authors:** Venkatesh Katari, Kesha Dalal, Narendra Kondapalli, Sailaja Paruchuri, Nagalakshmi Nadiminty, Charles K Thodeti

**Author notes:** Address all correspondence to: Charles Thodeti, Ph.D., Department of Physiology and Pharmacology, BHS 142; COMLS, The University of Toledo, 3000 Transverse Drive, Toledo 43614. Nagalakshmi Nadiminty, Ph.D., Department of Urology, The University of Toledo, 3000 Transverse Drive, Toledo 43614. This MS has been uploaded as a preprint at bioRxiv.

## Abstract

Bladder cancer (BLCA) is the second most common urologic cancer in the US and worldwide which mostly affects the aging population. Despite several ongoing clinical trials, treatment paradigms for BLCA have not changed significantly. Here, we investigated the expression of transient receptor potential vanilloid type 4 (TRPV4) in BLCA patients and its role in calcium influx, cell proliferation, and migration using normal human urothelial cells and BLCA cells. Bioinformatic analysis of the UALCAN and cBioPortal databases revealed that TRPV4 expression is significantly higher in human BLCA tissues compared to adjacent normal tissues. Further, the TRPV4 expression was markedly elevated in early-stage BLCA and upregulated in muscle-invasive bladder cancer (MIBC) tissues. TRPV4 is expressed in both normal urothelial (SV-HUC-1) and BLCA (T-24) cells and functional assays demonstrated enhanced TRPV4-mediated calcium influx in T-24 compared to SV-HUC-1 cells. T-24 cells exhibited higher spreading on extracellular matrix (ECM) gels with increasing stiffness (0.2, 8, and 50 kPa) and exhibited migratory phenotype compared to SV-HUC-1 cells. Pharmacological inhibition of TRPV4 significantly reduced proliferation and migration in T-24 cells but had minimal effects on normal cells. Finally, treatment with cisplatin significantly reduced TRPV4 protein levels and TRPV4-mediated calcium influx in chemosensitive UM-UC-3 cells, which remained unchanged in chemoresistant T-24 cells, suggesting a potential role of TRPV4 in chemoresistance. In conclusion, TRPV4 may contribute to bladder cancer progression by regulating cell proliferation and migration and may impart resistance to chemotherapy. Targeting TRPV4 could present a novel therapeutic approach for managing bladder cancer progression and overcoming chemoresistance.

**Significance Statement:** This study identifies TRPV4 as a critical driver of BLCA progression. Elevated TRPV4 gene expression in both early-stage and muscle-invasive bladder cancer (MIBC) tissues is associated with increased calcium signaling, cell proliferation, and migration. Importantly, TRPV4 inhibition selectively reduces BLCA growth and motility. Furthermore, TRPV4 is downregulated by cisplatin in chemo-sensitive but not chemo-resistant BLCA cells, underscoring its key role in bladder cancer chemoresistance. These findings position TRPV4 as a therapeutic target for enhancing BLCA treatment and overcoming drug resistance.

## 1. Introduction

Bladder cancer (BLCA) is the ninth most diagnosed cancer worldwide and the second most common urologic malignancy after prostate cancer, primarily affecting older adults (Bray et al., 2018; Saginala et al., 2020). Most BLCAs are urothelial in origin and are superficial, although about a third invade past the bladder submucosa/mucosa and are defined as muscle-invasive bladder cancer (MIBC) (Kaufman et al., 2009). Non-muscle invasive bladder cancer (NMIBC) is primarily treated with radical or partial cystectomy followed by adjuvant therapy. Up to 75% of NMIBC cases recur, necessitating long term surveillance; and as many as 25% progress to muscle-invasive disease, which is associated with a high risk of death from distant metastases (Witjes et al., 2014). Although several promising therapies such as immune checkpoint inhibitors, targeted therapies, and novel intravesical treatments are in development and clinical trials, standard treatment paradigms for BLCA have remained largely unchanged for decades, especially for patients who do not respond to initial therapies. Therefore, there is a need to understand the basic biology of the bladder cancer to identify new therapeutics targets.

Tumor growth and metastatic progression, characterized by epithelial-to-mesenchymal transition (EMT), invasion, migration, and angiogenesis, are regulated by changes in Ca^2+^ homoeostasis (Fels et al., 2018). One of the central players in regulating Ca^2+^ homoeostasis are plasma membrane-embedded calcium channels such as the transient receptor potential (TRP) superfamily of cation channels. TRPV4, a member of the TRP superfamily, is a non-selective cation channel that mediates calcium influx in response to biochemical and mechanical factors in many cell types including endothelial cells, epithelial cells, fibroblasts, and macrophages (Adapala et al., 2021). We have demonstrated that TRPV4 regulates tumor endothelial proliferation, migration, and tumor angiogenesis (Adapala et al., 2016; Thoppil et al., 2016). However, abnormal expression of TRPV4 has been linked to gastric, liver, pancreatic, colorectal, lung, and breast cancers (Yu et al., 2019; Yang and Kim, 2020; Zhang et al., 2021). A significant upregulation of TRPV4 was detected in metastatic breast cancer cell lines (Lee et al., 2016; Lee et al., 2017) and its expression increased with tumor grade and size and correlated with poor survival (So et al., 2020). TRPV4 has been shown to play an important role in normal bladder function (Gevaert et al., 2007). TRPV4 in the bladder urothelium contributes to sensory transduction, regulates barrier functions, modulates DSM function, and triggers the urethral to bladder reflex (Datta et al., 2021). The Inhibition of TRPV4 has been used to treat bladder dysfunction in mice and rats with cystitis (Everaerts et al., 2010). TRPV4 has been shown to be highly expressed in BLCA cells (Mizuno et al., 2014). These results implicate TRPV4 in the bladder dysfunction caused by several disease states including but not limited to cancer. However, the role of TRPV4 in bladder carcinogenesis and progression is relatively unknown. In the present study, we investigated the functional expression of TRPV4 in bladder cancer cells and its role in proliferation, migration, and chemoresistance. By targeting TRPV4, it might be possible to hinder the growth and spread of bladder cancer, potentially providing a new therapeutic option alongside existing treatments in the field.

## 2. Methods

### 2.1. UALCAN and cBioPortal databases

The University of Alabama at Birmingham Cancer Data Analysis Portal (UALCAN) is a comprehensive and user-friendly digital platform that permits in-depth analysis of cancer-related omics data. We have examined the differences of TRPV4 expression levels in bladder cancer tissue samples. We have retrieved TRPV4 expression from the patients (n=19) and bladder cancer primary tumor (n=408) (Chandrashekar et al., 2022) from the UALCAN database (https://ualcan.path.uab.edu/index.html). The cBioPortal for Cancer Genomics is a valuable online platform that enables the interactive exploration of genetic alterations and multidimensional cancer genomics data (Cerami et al., 2012). In the present study, we utilized the cBioPortal (https://www.cbioportal.org/) to investigate the genetic changes associated with the TRPV4 mRNA gene expression (Log2) across various stages (T0 to T4) of bladder cancer.

### 2.2. Cell culture and maintenance

Normal human uroepithelial cells (SV-HUC-1), human urothelial cancer cells (UM-UC-3T-24 and T-24) were purchased from the American Type Culture Collection (ATCC, USA). T-24 cells were cultured in McCoy’s 5A medium (Corning, USA) supplemented with 10% fetal bovine serum (FBS). UM-UC-3 cells were cultured in EMEM supplemented with 10% FBS. SV-HUC-1 cells were cultured in F-12 K medium (Corning, USA) with 10% FBS. All cells were grown in plastic tissue culture T-flasks (Corning, USA) at 37 °C and 5% CO_2_. Cell lines were authenticated by American Type Culture Collection, USA.

### 2.3. Calcium influx assay

Cells cultured on MatTek glass-bottomed dishes or cover glasses were loaded with Fluo 4-AM (4 μM) for 30 min, washed three times in calcium medium (in mM: 136 NaCl, 4.7 KCl, 1.2 MgSO4, 1.1 CaCl2, 1.2 KH2PO4, 5 NaHCO3, 5.5 glucose, and 20 HEPES, pH 7.4), and kept in this medium on an inverted Leica confocal microscope (SP5). Cells were stimulated with GSK1016970A (100 nM), and images were acquired every 3 s and analyzed using Leica software and Microsoft Excel as previously described (21, 33). In some experiments, cells were pretreated with cisplatin (1 μM) for 72 h before measuring calcium influx.

### 2.4. Cell proliferation assay

Cell viability was evaluated using the TACS XTT Cell proliferation kit, based on the 2,3-Bis-(2-Methoxy-4-Nitro-5-Sulfophenyl)-2H-Tetrazolium-5-Carboxanilide (XTT), according to the protocol described by the manufacturer (R&D Systems, USA). Briefly, 2 x 10^3^ cells were plated in each well of a 96-well plate and incubated at 37 C° in a 5% CO_2_ atmosphere saturated with water vapor for 24 h; they were then incubated for 24 h in medium containing with or without GSK2193874 (GSK2; 50 nM). Cellular viability in the absence and presence of GSK2 was quantitated calorimetrically by examining absorbance at 450 nm (A_450_). Control cells were treated in the same way, but without GSK2 and the relative cell viability was determined.

### 2.5. Transwell cell migration assay

The transwell migration experiment was performed with the 8 μm-pore chambers inserted into 24-well plates (Corning, NY, USA). Both SV-HUC-1 and T-24 cells were plated in the upper chambers and 600 μL DMEM containing 30% FBS was placed at the bottom of the chamber. After 24 h incubation, cells were fixed with 4% paraformaldehyde in PBS and stained with crystal violet for 20 min. The images of cells that migrated to the lower surface of the membrane were acquired with an inverted microscope, and the number of migrated cells was counted.

### 2.6. Western blot analysis

Cells were lysed in ice-cold Triton-X 100 buffer, and protein concentration was determined by a BCA protein assay kit (Santa Cruz, CA, USA). Cell lysates were mixed with reducing sample buffer, boiled for 10min and subjected to SDS-PAGE and electroblotted onto PVDF membranes. Membranes were incubated sequentially with TRPV4 primary antibodies (1:300 Alomone, USA) and anti-rabbit (1:5000) secondary antibodies and developed using a western blot imaging system (Azure biosystems, USA). The band intensities were quantified using Image J.

### 2.7. Immunofluorescence staining

Cells were fixed in 4% paraformaldehyde in phosphate buffer at pH 7.4. Cells were incubated at RT for 20 minutes and washed three times with PBS. Cells were then permeabilized with 0.1% Tritonx-100 for 15 minutes and washed three times with PBS. Cells were blocked with complete media containing 10% FBS for 30 min. Finally, cells were incubated with primary antibodies vinculin (1:100) and F-actin (1:100) for 1 h at RT. Cells were washed three times with PBS and incubated with secondary antibodies (anti-rabbit or anti-mouse Alexa 488 or 594) for 1 hour at RT in a dark room. After the incubation cells were washed and mounted on slides using Fluormount containing DAPI (Thermofischer Scientific, USA) and images were acquired on Leica SP5 confocal laser scanning microscope.

### 2.8. Mouse xenograft assays

Male 4–5-week-old SCID mice (Charles River, Wilmington, MA) with an average weight of 20 g were allowed to acclimate for 7 days after receipt from the vendor and were housed at 22.5 ± 0.5°C in sterile cages. UM-UC-3 cells (2x10^6^/flank) were injected along with 1:1 matrigel sub-cutaneously into both flanks of mice and tumor growth was monitored. When the tumor volumes reached approximately 0.1 cm^3^, mice were randomly divided into groups (n=5/group) and treated with: 1) vehicle control (0.5% Methocel A4M) and 2) 0.5 mg/kg cisplatin i.p. every other day. Treatments were performed for 3 weeks, and tumor growth (using digital calipers) and mouse weights (using digital balance) were monitored every other day. Mice were euthanized with carbon dioxide followed by cervical dislocation. Tumor tissues were harvested and the expression level of TRPV4 was analyzed using western blotting. All animal experiments were performed after approval from the Institutional Animal Care and Use Committee of the University of Toledo (IACUC protocol # 108804) and in accordance with the National Institutes of Health guide for the care and use of laboratory animals.

### 2.9. Statistical Analysis

The data are shown as mean ± SE. Statistical significance was determined using analysis of variance with Bonferroni’s comparison test. The Welch’s T-test was used in all studies performed by the UALCAN database. Significant differences are indicated in the figures such as \**p≤0.05* or \*\**p<u><0</u>.01*. \*\*\**p<u><0</u>.001*, \*\*\*\**p<u><0</u>.0001*.

## 3. Results

### 3.1. TRPV4 expression is higher in advanced BLCA tissues

To investigate the role of TRPV4 in BLCA, we analyzed TRPV4 mRNA expression in BLCA clinical samples from the UALCAN and The Cancer Genome Atlas Program (TCGA) datasets using cBioPortal databases. We found that primary bladder cancer tissues express significantly higher levels of TRPV4 compared to normal tissues (Fig. 1A). Next, we measured TRPV4 transcript levels by qPCR in bladder cancer tissues from the University of Toledo Urology Biospecimen Biorepository. Our results showed that TRPV4 mRNA levels are elevated in Muscle Invasive Bladder Cancer (MIBC) tissues (Fig. 1B). Importantly, we observed that TRPV4 expression is higher in malignant bladder cancer cells than in immune cells (monocytes, macrophages, and dendritic cells) within the tumor microenvironment (Fig. 1C). These findings suggest that TRPV4 may play a role in promoting bladder tumorigenesis.

**Figure 1.**
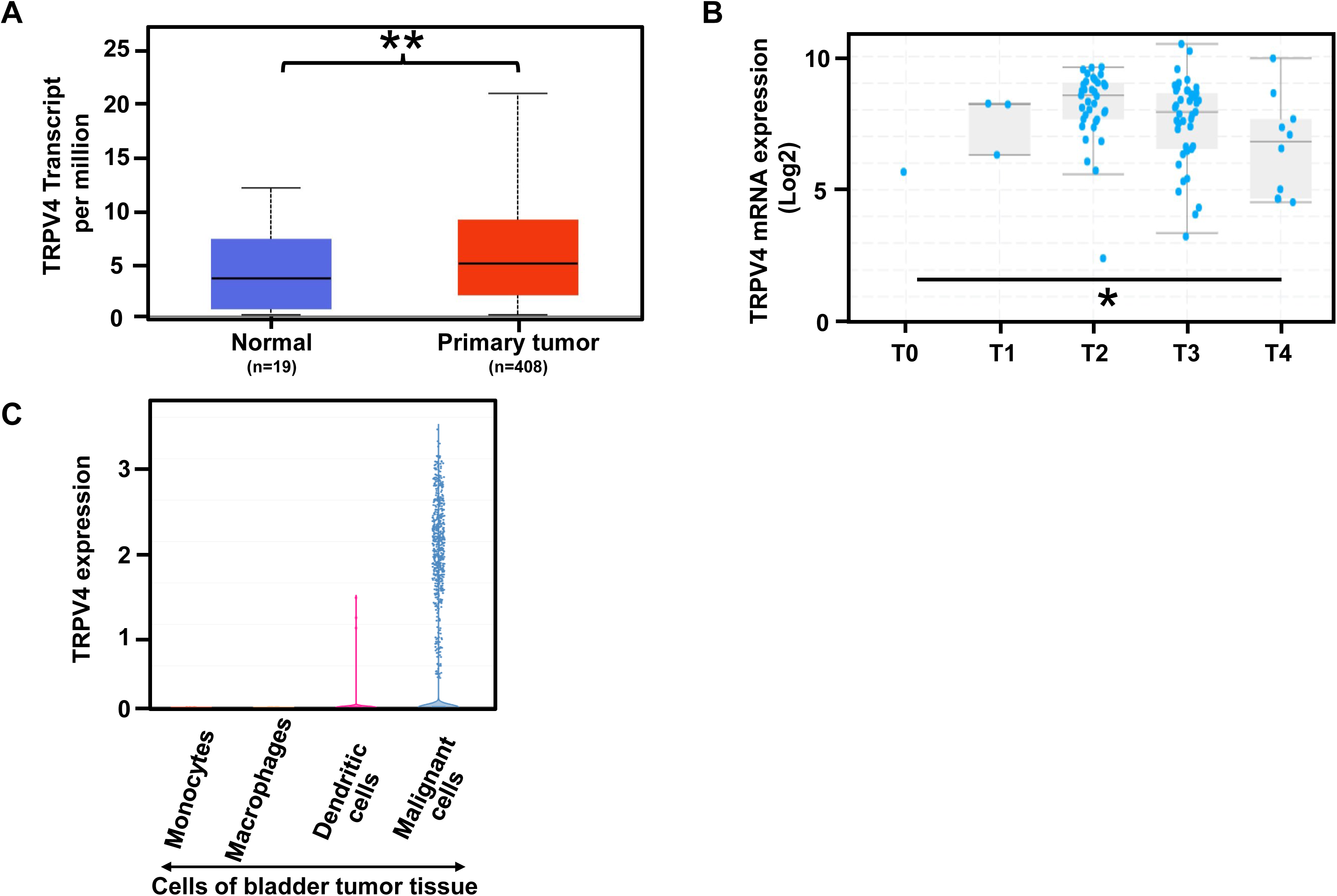
TRPV4 expression in bladder cancer tissues. **A)** TRPV4 transcript expression in human bladder tissues (Normal n=19; tumor n=408; *p ≤ 0.01*). **B)** Bladder samples from the MSKCC, TCGA datasets demonstrated a significant (*p≤0.05*) upregulation of TRPV4 mRNA levels during disease progression. Data normalized from Illumina Hiseq-RNA seq (log2). **C)** TRPV4 expression in malignant cells and immune cells (monocytes, macrophages, dendritic cells) in bladder tumor tissues.

### 3.2. TRPV4 is functionally expressed in BLCA cells and is correlated with BLCA aggressiveness

Next, to determine the role of TRPV4 in BLCA, we confirmed the functional expression of TRPV4 in BLCA cells (UM-UC-3, HT-1197, and T-24) and normal urothelial cells (SV-HUC-1) using western blotting, and calcium imaging (Fig. 2 and Fig. 5). We found that TRPV4 is expressed at higher levels in the higher-grade T-24 cells compared with normal SV-HUC-1 cells (Fig.2A). UM-UC-3 cells expressed similar levels of TRPV4 to that of T-24 (Fig. 5A). Importantly, a specific agonist of TRPV4, GSK1016790A (GSK), induced robust calcium influx in BLCA cells (Fig. 2B). Notably, we found that TRPV4-induced calcium influx is higher in T-24 cells compared with normal urothelial cells (Fig. 2B, C) suggesting that TRPV4 functional expression may correlate with the tumorigenic potential of BLCA cells.

**Figure 2.**
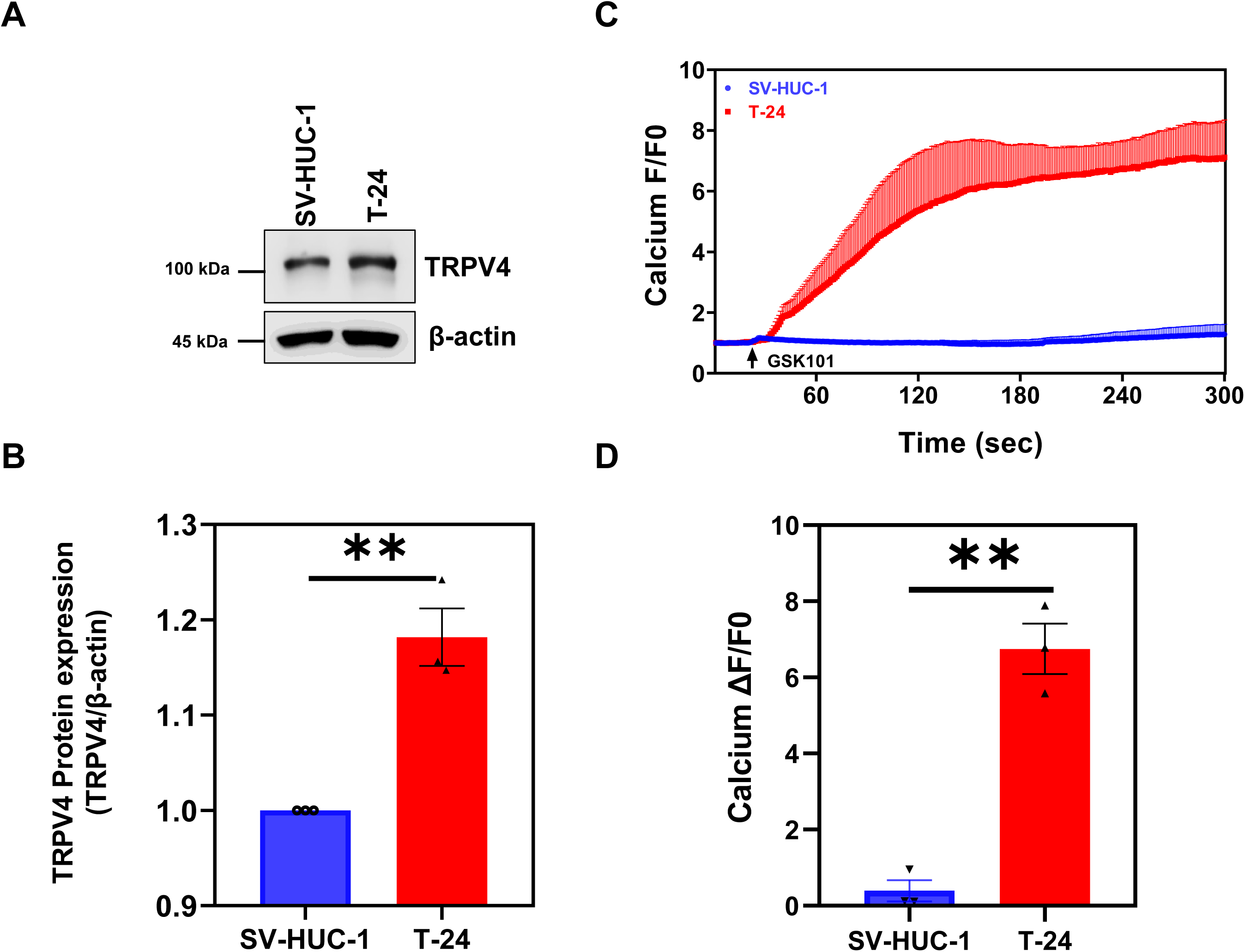
TRPV4 functional expression in bladder cancer cells. **A)** Western blot showing TRPV4 expression in SV-HUC-1 and T-24 BCLA cells. Note: increased TRPV4 protein in bladder cancer (T-24) cells compared to normal cells (SV-HUC-1) which is correlated with increased calcium influx. **B)** Quantification of western blot bands of TRPV4 normalized to β-actin**. C)** Transients showing calcium influx in response to TRPV4 agonist GSK1016709A (GSK101) in Fluo-4 loaded T-24 and SV-HUC-1 cells. **D)** Quantification shows higher TRPV4-mediated calcium influx in T-24 cells compared to SV-HUC-1 cells. The data shown is mean ± SEM from three independent experiments. Significance was determined using Student’s paired t-test (\*\**p ≤ 0.01*).

### 3.3. TRPV4 is required for matrix stiffness-mediated BLCA cell attachment and spreading

To investigate whether functional expression of TRPV4 is critical for matrix stiffness-mediated BLCA cell spreading, we cultured these cells on ECM gels with increasing stiffness levels (0.2, 8, and 50 kPa) that mimic the range from normal tissue to tumor stiffness *in vivo*. We observed that T-24 BLCA cells exhibited significantly higher spreading across all stiffness levels compared to normal SV-HUC-1 cells (Fig. 3A), indicating enhanced mechanosensing mediated by TRPV4. Additionally, immunofluorescence staining revealed prominent stress fibers and focal adhesions in T-24 cells, which were notably distinct from those in normal SV-HUC-1 urothelial cells (Fig. 3B).

**Figure 3.**
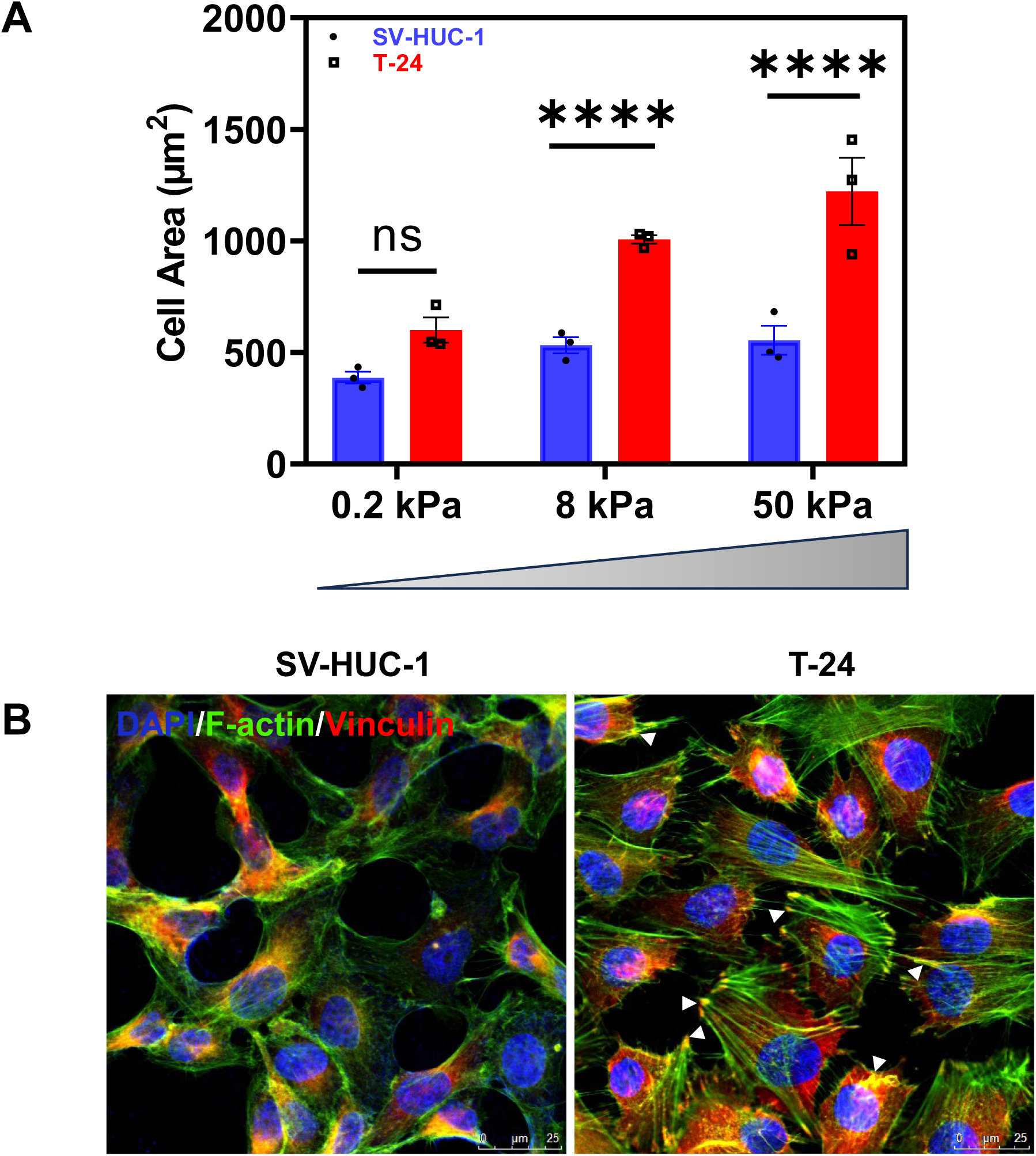
TRPV4 mechanosensitivity towards ECM stiffness in human bladder cancer cells. **A)** ECM stiffness-dependent spreading of SV-HUC-1 and T-24 cells cultured on ECM gels that mimic stiffness of the tumor (0.2, 8 and 50 KPa). Quantification of cell spreading (cell area) shows increased spreading of T-24 cells with the increase in ECM stiffness compared to SV-HUC-1 cells. The data shown is mean ± SEM from three independent experiments. Significance was determined using Two-way ANOVA (\*\*\*\**p ≤ 0.0001; ns = p > 0.05*). **B)** Representative confocal microscopic images showing stress fibers (F-actin:green) and focal adhesions (vinculin=red) in normal (SV) and T24 cells (n=3 independent experiments). Note colocalization of stress fibers with focal adhesions (arrows) (Scale bar: 25µm).

### 3.4. BLCA cells exhibit TRPV4-dependent proliferation and migration

We next assessed proliferation and migration of key hallmarks of metastatic cancer in T-24 and SV-HUC-1 cells treated with or without the specific TRPV4 antagonist, GSK2. XTT assays showed that T-24 cells proliferate significantly more than SV-HUC-1 cells (Fig. 4A). Treatment with GSK2 markedly reduced proliferation in T-24 cells, while having no effect on SV-HUC-1 cells (Fig. 4A). Additionally, transwell migration assays demonstrated that T-24 cells migrated more extensively than SV-HUC-1 cells, as indicated by the increased number of migrating cells (Fig. 4B). This enhanced migration was inhibited by GSK2 treatment. Quantitative analysis confirmed significantly higher migration in T-24 cells, with GSK2 reducing migration levels to those comparable with SV-HUC-1 cells (Fig. 4C).

**Figure 4.**
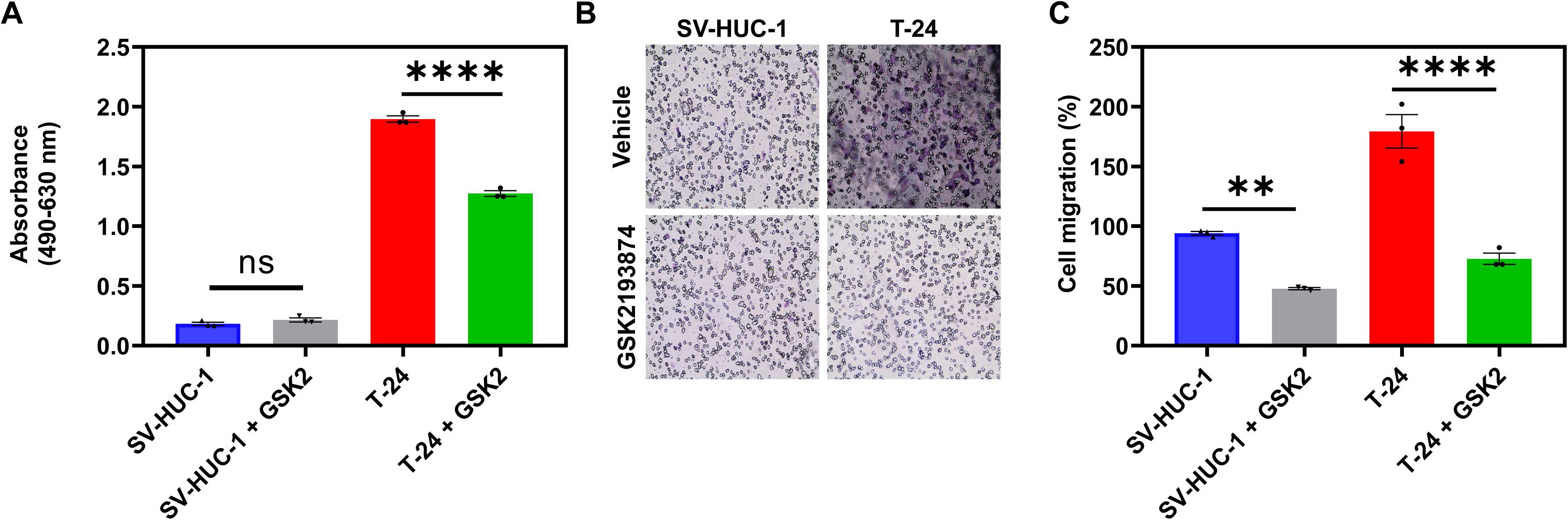
TRPV4 regulates bladder cancer cell proliferation and migration. **A)** XTT assay showed higher proliferation in T-24 BCLA cells compared to SV-HUC-1 cells, which was significantly reduced by GSK2 (100 nM). ns = *p > 0.05*. **B)** Representative Brightfield crystal violet stained images (4×) of Transwell migration assay showing higher migration (blue) of T-24 cells compared to SV-HUC-1 cells. **C)** Quantitative analysis showing higher migration in T-24 cells compared to SV-HUC-1 cells. Pharmacological inhibition of TRPV4 with GSK2 attenuated higher migration exhibited by T-24 BCLA cells. The data shown is mean ± SEM from three independent experiments. Significance was determined using Two-way ANOVA and significance was set at ** *p<0.01* and \*\*\*\**p ≤ 0.0001*.

### 3.5. TRPV4 imparts resistance to chemotherapy in BLCA cells

The preceding results suggest that functional expression of TRPV4 may influence the tumorigenic potential of BLCA cells, as reflected by their proliferation and migration. To investigate whether TRPV4 also affects the chemosensitivity of BLCA cells, we treated UM-UC-3 (cisplatin-sensitive) and T-24 (cisplatin-resistant) cells with 1 µM cisplatin for 72 hours in vitro and measured TRPV4 expression by western blotting. As shown in Fig. 5A, cisplatin treatment significantly downregulated TRPV4 expression in UM-UC-3 cells but had no effect on T-24 cells. Moreover, GSK1-induced (TRPV4-mediated) calcium influx was significantly reduced in cisplatin-treated UM-UC-3 cells compared to untreated controls, whereas no difference in calcium influx was observed between untreated and cisplatin-treated T-24 cells (Fig. 5B). Finally, TRPV4 expression was reduced in cisplatin-treated UM-UC-3 xenografts relative to vehicle-treated controls (Fig. 5C).

**Figure 5.**
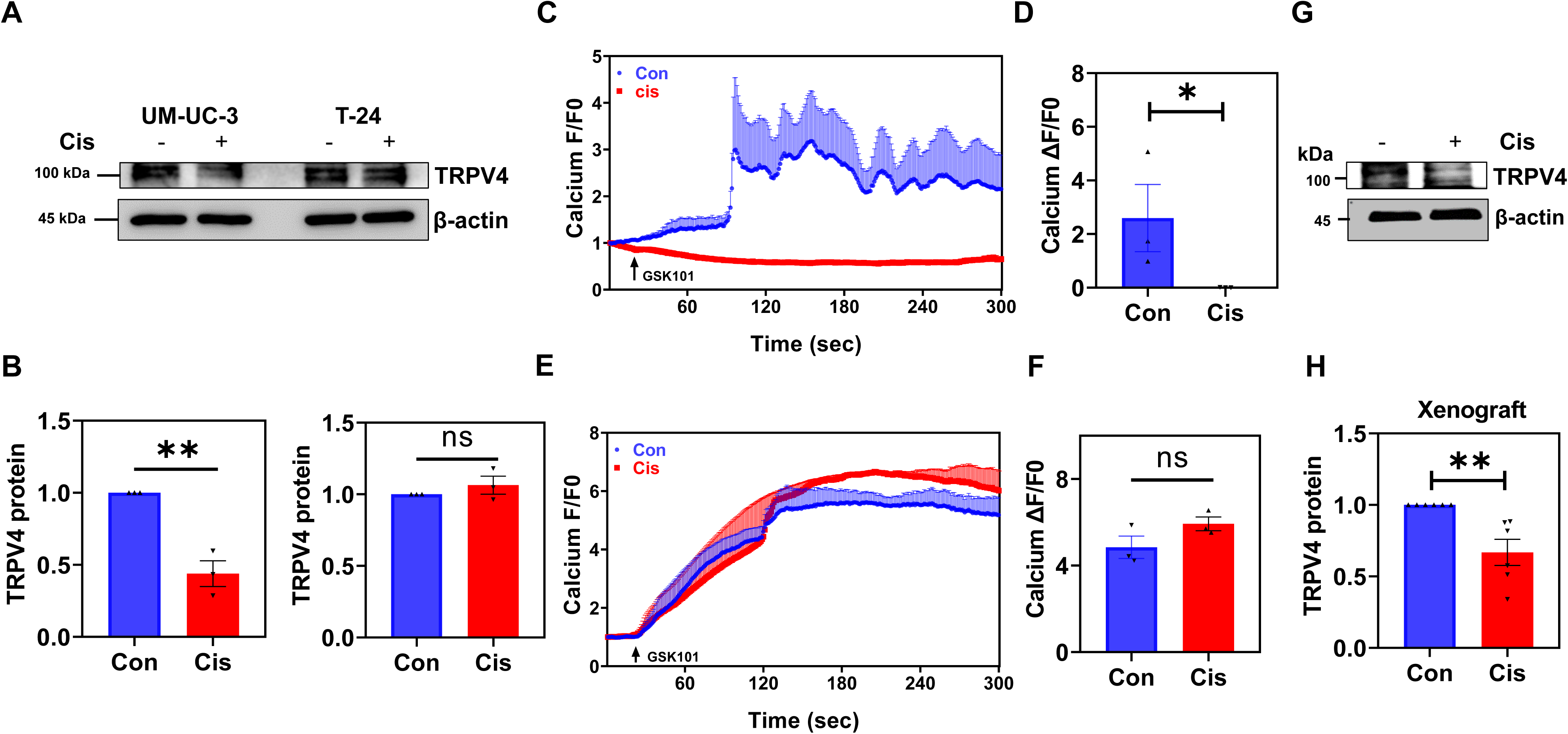
TRPV4 may impart resistance to Cisplatin therapy in bladder cancer. **A)** Western blot showing cisplatin-induced downregulation of TRPV4 in chemoresponsive UM-UC-3 cells compared to resistant T-24 cells in vitro. **B)** Quantification of the western blots demonstrate significant down regulation of TRPV4 in UM-UC-3 cells compared to T-24 cells and the band intensities were normalized by β-actin (n=3, \*\**p≤ 0.01*). Transients showing calcium influx in response to TRPV4 agonist GSK1016709A in Fluo-4 loaded UM-UC-3 (**C**) and T-24 (**E**) cells. Quantification shows higher TRPV4-mediated calcium influx in UM-UC-3 (**D**) and T-24 (**F**) cells before and after treatment with Cisplatin. **G)** Western blot showing TRPV4 expression in UM-UC-3 xenografts in mice treated with vehicle or Cisplatin (Cis). **H)** Quantification of the western blots demonstrates significant down regulation of TRPV4 in UM-UC-3 cells in response to Cisplatin. The data shown is mean ± SEM from three independent experiments. Significance was determined using Student’s paired t-test and significance was set at \*\**p<0.01* and ns = p>0.05.

## 4. DISCUSSION

In this study, we provide evidence for the critical role of TRPV4 in bladder cancer progression and chemoresistance. This conclusion is supported by several key findings:

1) TRPV4 expression is elevated in bladder cancer tissues compared to normal tissues and correlates with tumor aggressiveness; 2) TRPV4 is functionally expressed at higher levels in high-grade, invasive T-24 cells; 3) T-24 cells with high TRPV4 expression exhibit enhanced proliferation and migration relative to normal urothelial cells, both of which are attenuated by the specific TRPV4 antagonist GSK2; and 4) TRPV4 expression remains unchanged in chemoresistant T-24 cells but is reduced in chemosensitive UM-UC-3 cells following cisplatin treatment.

TRPV4 is known to function as a mechanically activated channel in various cell types including endothelial, fibroblast, and renal epithelial cells (Thodeti et al., 2009; Adapala et al., 2020; Adapala et al., 2023; Kondapalli et al., 2023) and has been implicated in several cancers such as breast, gastric, and prostate cancers. However, its role in bladder cancer has not been characterized. Through analysis of clinical datasets and experimental validation, we established that TRPV4 expression is elevated in MIBC tissues and in higher-grade BLCA cells. The increased TRPV4 mRNA levels observed across different bladder cancer stages suggest a mechanistic role in tumor progression.

Consistent with this, our calcium influx assays demonstrated higher functional TRPV4 expression in bladder cancer T-24 cells compared to normal urothelial SV-HUC-1 cells, implicating TRPV4 activity in the tumorigenic phenotype. Mechanosensing is crucial for tumor progression, enabling cancer cells to adapt and interact with their microenvironment. We found that bladder cancer cells display increased spreading on ECM gels of increasing stiffness, with a strong correlation between TRPV4 expression and mechanosensitivity. The increased projected cell area of T-24 cells on stiffer ECM substrates (50 ≥ 8 ≥ 0.2 kPa), combined with their enhanced migration, underscores the role of TRPV4 in sensing and responding to biomechanical cues. These findings suggest that TRPV4-mediated mechanotransduction may play a vital role in bladder cancer cell adhesion, invasion, and potentially metastasis.

Proliferation and migration are essential hallmarks of tumor progression. Our study demonstrates that the inhibition of TRPV4 using GSK2193874 significantly reduced proliferation and migration in bladder cancer cells, indicating that TRPV4 is a key driver of bladder cancer progression. Moreover, TRPV4 appears to regulate bladder cancer cell migration through modulation of cytoskeletal components such as vinculin and F-actin. The distinct presence of stress fibers and focal adhesions in T-24 cells which were absent or reduced in normal urothelial cells supports the hypothesis that TRPV4 enhances migratory and invasive potential. This is consistent with prior reports on other cancers, where TRPV4 promotes migration via cytoskeletal remodeling (Lee et al., 2016; Huang et al., 2022; Pu et al., 2022; Ren et al., 2025).

Our findings also provide novel insights into the role of TRPV4 in bladder cancer chemoresistance. We observed that TRPV4 expression remained stable in cisplatin-resistant T-24 cells after treatment but was significantly downregulated in cisplatin-sensitive UM-UC-3 cells. Calcium influx assays confirmed reduced TRPV4-mediated calcium influx in cisplatin-treated UM-UC-3 cells but no change in T-24 cells, suggesting that TRPV4 may contribute to chemoresistance by sustaining intracellular signaling pathways that promote survival under chemotherapeutic stress. Additionally, reduced TRPV4 expression in cisplatin-treated UM-UC-3 xenografts supports the idea that TRPV4 expression levels may influence therapeutic response. Together, these results highlight TRPV4 as a potential biomarker for chemoresistance and a promising therapeutic target for improving bladder cancer treatment outcomes.

Interestingly, the importance of TRPV4 appears context-dependent across cancer types. While TRPV4 activation induces apoptosis in breast cancer cells, it promotes proliferation and migration in gastric cancer. Our study aligns with the latter, showing that TRPV4 inhibition attenuates BLCA proliferation and migration, indicating that TRPV4 activation supports bladder cancer progression. However, the molecular mechanisms regulating TRPV4 activation in cancer remain unclear. Given that bladder cancer predominantly affects older populations (Taylor and Kuchel, 2009), it is noteworthy that TRPV4 expression increases with age in human and mouse hearts, where it has been implicated in calcium handling dysregulation and cardiac dysfunction (Jones et al., 2019; Peana et al., 2022). Similarly, elevated functional TRPV4 expression may activate tumorigenic pathways in bladder cancer; a hypothesis warranting future investigation.

## 5. Conclusions

In summary, our study provides compelling evidence that TRPV4 is upregulated in advanced bladder cancer and plays a crucial role in mechanosensing of the tumor matrix, tumor progression, and chemoresistance. The functional link between TRPV4 activity and bladder cancer aggressiveness suggests that targeting TRPV4 could represent a novel therapeutic approach. Future research should aim to elucidate the precise molecular mechanisms by which TRPV4 modulates tumorigenic signaling pathways and explore TRPV4 inhibitors as potential therapeutic agents in bladder cancer management.

## Abbreviations

BLCA, Bladder cancer; TRPV4, Transient receptor potential vanilloid type 4; ECM, Extracellular matrix; NMIBC, Non-muscle invasive bladder cancer; EMT, Epithelial-to-mesenchymal transition; UALCAN, The University of ALabama at Birmingham CANcer data analysis Portal.

## Data Availability Statement

The data to support the findings of the present study are available from the corresponding author upon reasonable request.

## Author Contributions

*Participated in research design:* Venkatesh Katari, Nagalakshmi Nadiminty, Charles K Thodeti

*Conducted experiments:* Venkatesh Katari, Nagalakshmi Nadiminty, Kesha Dalal, Narendra Kondapalli

*Analyzed and interpreted the results*: Venkatesh Katari, Nagalakshmi Nadiminty, Charles K Thodeti, Sailaja Paruchuri

*Writing of the manuscript*: Venkatesh Katari, Sailaja Paruchuri, Nagalakshmi Nadiminty, Charles K Thodeti

